# A Unified Neural Timecourse for Words, Phrases, and Sentences: MEG Evidence from Parallel Presentation

**DOI:** 10.1101/2025.11.17.688866

**Authors:** Nigel Flower, Liina Pylkkänen

## Abstract

Recent behavioral and neural research on reading shows that humans can extract syntactic structure from short sentences within a fraction of a second—faster than many estimates for recognizing the meaning of a single word. This challenges a core assumption of many language processing models: that combinatory operations depend on prior lexical access. Further, studies using parallel presentation of full sentences have revealed electrophysiological responses remarkably similar to those well established for single words. This raises the question of whether words, phrases, and sentences all move through the same processing stages, regardless of syntactic complexity. Using magnetoencephalography (MEG), we examined how single words, phrases, and sentences are processed when all visual information is available at once. Across all three levels, we observed highly similar waveform dynamics, with early responses reflecting bottom-up detection of form followed by activity in the left anterior and posterior temporal cortices and vmPFC consistent with combinatory processing. Of these regions, the left anterior temporal lobe (LATL) showed effects of bigram frequency suggestive of serial left-to-right dynamics. Together, these results support a Global-to-Serial Assembly (GLOSA) model in which the brain first detects the global form of the stimulus in a snapshot-like manner and then probes its combinatory properties through partially serial processes.

## INTRODUCTION

When listening to serially unfolding spoken language, it is natural for comprehension to occur incrementally word-by-word, as predicted by many psycholinguistic theories of sentence processing (Hale, 2001; Lewis & Vasishth, 2005). But what if this seriality was mainly a reflection of how the speech signal incrementally arrives into our perceptual system, as opposed to an inherent property of the language system? Recent results suggest that, in striking contrast to auditorily presented language, glancing at a written sentence may allow the brain to begin extracting basic phrase-structure information as early as 130 ms after presentation onset (Fallon & Pylkkänen, 2024), resembling how we rapidly grasp the gist of a visual scene (Wiesmann & Võ, 2022). Further, well-formed sentences perceived at-a-glance such as *the man can run* diverge from their scrambled counterparts (*run man can the*) in the N400 ERP component (Wen et al. 2019, 2021), a signal thought to reflect lexical processing (Lau et al., 2008). Both results challenge the view that language processing proceeds serially in all circumstances, including when the whole linguistic stimulus is available to the brain’s perceptual systems at the same time as in the case of reading.

The effects reported in Fallon and Pylkkänen (2024) and Wen et al. (2019, 2021), all using the psycholinguistic technique of rapid parallel visual presentation or RPVP, suggest that neural responses to full sentences may not be qualitatively different from those of single words. The similarity between the brain’s basic evoked response to a visually presented full sentence and its well-studied response to a single word is visually obvious—though not much discussed—in existing EEG studies: sentences elicit the same P1–N2–N400 ERP sequence observed for single words (e.g., Fig. 2 in Wen et al., 2021). Could it be, then, that when the brain sees a full sentence, it executes the same basic sequence of comprehension operations as it does for a single word, detecting form, structure and meaning? And if this is the case, to what extent is the processing at each of these stages characterized by serial or parallel mechanisms?

To reiterate, our first question is whether evoked responses to entire sentences resemble those elicited by single words, as suggested by prior work. One possibility is that combinatorial processing for sentences presented in parallel differs fundamentally from that of isolated words. On this *fully serial* account, combinatorial processing unfolds as described in canonical psycholinguistic models: lexical representations are incrementally accessed and integrated into an evolving syntactic and semantic structure (Frazier and Fodor, 1978; Frazier 1987; Hale, 2001; Lewis and Vashishth, 2005). This hypothesis predicts that sentence-evoked responses should differ qualitatively from those elicited by individual words. However, findings from Wen et al. (2019; 2021) and Fallon & Pylkkänen (2024) argue against this view. These studies show that combinatory effects for multi-word expressions presented in parallel emerge as quickly—or even more quickly—than those for serial input. If composition depended on lexical access, the opposite pattern would be expected, since multiple lexical items would need to be accessed before combination. Moreover, the waveforms for sentences and single words appear visually similar. Still, because these studies do not independently estimate the timing of lexical access, the serial hypothesis cannot yet be conclusively ruled out.

Under a less extreme version of the serial hypothesis, combinatorial processing proceeds incrementally, word-by-word, *after* form-based information has been fully encoded. In this account, the brain first takes a “snapshot” of the sentence and then builds a sentential representation from left to right. We term this the Global-to-Sequential Assembly (GLOSA) hypothesis. It assumes that sentence comprehension remains serial in its internal combinatorial mechanism but diverges from canonical models by positing that processing a sentence presented in parallel begins with a global, bottom-up encoding stage akin to that for a single word. Neural activity should therefore reflect two different stages: (1) an initial phase of perceptual encoding, followed by (2) combinatory processing in which each word is successively integrated into a sentential structure. Specifically, GLOSA predicts that encoding-related activity will precede combinatory effects, and that these combinatory effects will emerge earlier for words appearing earlier in the sentence and later for those appearing later.

A second possibility is that after taking a global snapshot of the stimulus, the brain integrates lexical information into a sentential representation in a single, parallel step. We term this the Global-to-Parallel Assembly (GLOPA) hypothesis. Like GLOSA, it posits an initial stage of bottom-up encoding followed by combinatorial processing, implying a shared early response profile for words and sentences. However, GLOPA proposes that combinatorial processing occurs simultaneously across words once the stimulus is fully encoded. It thus shares GLOSA’s prediction that perceptual encoding precedes combination but further predicts that combinatory effects for different bigrams will overlap in time.

At the extreme parallel end, combinatorial processing may unfold in a single, uninterrupted step. We term this the Global Unified Assembly (GLU) processing hypothesis. Unlike GLOSA and GLOPA, GLU posits that form-based encoding and combinatory processing unfold concurrently from the very onset of perception, producing a sentential representation in one continuous neural process. As a result, sentence-evoked responses are predicted to be fundamentally distinct from those elicited by individual words: the neural signal never reflects isolated word-level processing, but instead represents integrated sentence-level structure from the first moment. Supporting this possibility, Fallon & Pylkkänen (2024) report sensitivity to syntactic phrase structure as early as 125 ms after sentence onset, suggesting that the brain can detect structural cues extremely rapidly. These early effects make it plausible that combinatorial processing is ongoing from the outset of perception.

The hypotheses outlined above differ in the timing and degree of parallelism that they assume, but direct neural evidence is limited. While behavioral and neuroimaging studies have begun to illuminate the brain’s capacity for at-a-glance sentence processing, it remains unclear how sentence-level responses relate to those evoked by single words. Initial RPVP studies have provided a window into this question: Flower and Pylkkänen (2024) used MEG to show a limited degree of parallel processing in short four-word sentences, and electroencephalography (EEG) signals reveal overlapping sensitivity to the third, fourth, and fifth words of five-word sentences within 160-300ms.

Although prior work has characterized combinatorial processing for individual words and short phrases using serial presentation (Pylkkänen, 2019), little is known about the neural signatures elicited when the brain perceives an entire sentence at once. To address this gap, we designed an MEG experiment manipulating both the size and compositionality of stimuli, allowing us to test the predictions of GLOSA, GLOPA, and GLU. Compositional conditions involved full sentences (e.g., the cats are nice), determiner phrases (the cats) and bare determiners (the). Non-combinatory controls were lists of one, two, or three plural nouns, blocking noun-noun composition. We first conducted a region of interest (ROI) analysis targeting cortical areas previously implicated in MEG studies of combinatorial processing. The goal was to identify two types of effects: size effects, reflected as increased neural activity for larger stimuli regardless of compositionality, and combinatory effects, reflected as increased neural activity for compositional conditions only. GLOSA and GLOPA predict that size effects should precede combinatory effects, reflecting the idea of first taking a “snapshot” of the stimulus and then computing its compositional meaning. In contrast, GLU predicts size and combinatory effects to emerge concurrently, consistent with a single-stage, fully parallel comprehension mechanism.

We then applied a two-stage generalized linear model (GLM) analysis within the significant clusters identified in the ROI analysis to characterize their computational properties. We operationalized lexical access as a region’s sensitivity to lexical frequency and compositional processing as its sensitivity to bigram frequency. This analysis tested whether combinatory effects followed a serial or parallel profile, with serial effects being compatible with GLOSA and parallel effects being compatible with GLOPA or GLU.

## MATERIALS AND METHODS

### Participants

Twenty-five right-handed speakers of English with normal or corrected-to-normal vision participated in this study. Each participant gave informed consent. One recording was excluded from analysis due to excessive sleepiness, resulting in a total of twenty-four participants for the behavioral and MEG analyses.

### Stimuli

The principal aim of this study was to understand the temporal dynamics of neural activity in response to sentences versus single words or small phrases. Thus, the experiment consisted of stimuli ranging from single words, determiner phrases, and full sentences. The sentences were formed by collecting 50 adjective-noun pairs such as *cats*-*nice* and inserting one of four determiners in front of the noun and a plural copula between the noun and the adjective, yielding sentences such *as all cats are nice*. The determiner phrases were formed by removing the copulas and adjectives from the sentence stimuli, and the bare determiners simply consisted of the determiners *all*, *some*, *no*, or *the*. The explicit manipulation of the determiner was intended to investigate matters separate from this current project. This yielded a 3 × 4 design (Fig. 2) with the factors Size (Bare Determiner, Determiner Phrase, Sentence) and Determiner (all, some, no, the). A second goal of this study was to assess whether the neural activity we see for sentences compared to words or phrases is specific to compositional processing as opposed to a mere effect of stimulus size. To address this, we included non-compositional conditions, which consisted of the plural nouns used in the compositional stimuli organized into sequences of one, two, or three nouns. Plural nouns were chosen specifically for the non-compositional stimuli, because they could not be composed into noun-noun compounds. When the combinatory stimuli showed effects of size that did not replicate for the non-combinatory stimuli, we deemed such signals to reflect composition, not mere stimulus size.

When presenting stimuli containing multiple words in parallel, it was important for us to ensure that participants could comprehend their content within a single glance, similar to one’s experience of, say, glancing at a notification on one’s phone. We thus explicitly controlled the number of characters that each noun and adjective such that the full sentences and the noun lists would not stretch beyond the parafovea, 8 degrees from the visual field. Nouns were restricted to be no longer than three characters long, resulting in plural forms no longer than four characters. Adjectives varied between three and four characters in length. The sentence conditions contained the largest stimuli and had an average length of 16.72 characters with a standard deviation of 0.84, with a range of 15 to 18 characters. Lastly, the sentence conditions occupied on average 6.11 degrees of the visual field, ensuring that the largest of the stimuli in this experiment were squarely within the parafovea.

**Figure 1:**
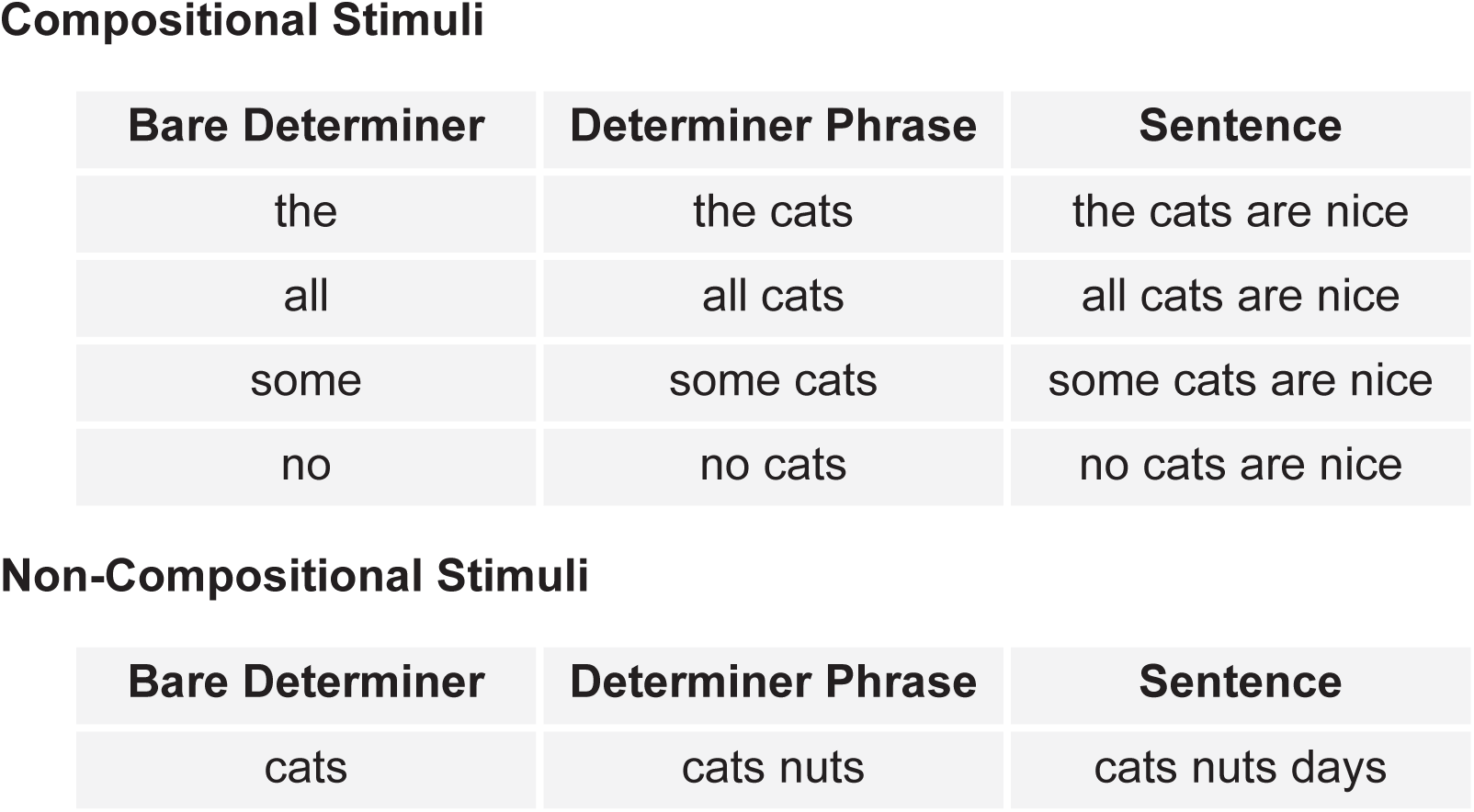
Factorial design of experiment. The stimuli in the top box enumerate compositional stimuli that vary in size as well as determiner. Bottom box lists the non-compositional control stimuli ranging from a single plural noun to three plural nouns.

### Trial structure and task

Trials began with a fixation cross onscreen for 200 ms followed by a blank screen for another 200 ms. Next, a stimulus was presented for 300 ms followed by a blank screen for 500 ms. After this presentation, a second stimulus appeared onscreen that was either identical to the first stimulus or differed from the first stimulus by replacing one word with another word of the same syntactic category drawn from the set of words used in creating the stimuli. This second stimulus remained onscreen until subjects responded whether it was a “match” or a “mismatch” with the first stimulus. The matching task was identical to that of Fallon & Pylkkänen (2024) and Flower & Pylkkänen (2024), which used an adaptation of the matching tasks in Pegado and Grainger (2020). The benefits of the task include its simplicity and the fact it could in principle be accomplished through purely perceptual mechanisms, and thus, any effects of higher-level stimulus factors necessarily reflect automatic language processing, as opposed to any type of metalinguistic behavior. Prior studies have reported clear Sentence Superiority Effects with this task, i.e., faster and more accurate responses for sentences than for unstructured stimuli, indicating a rapid deployment of grammatical knowledge (Fallon & Pylkkänen, 2024; Flower & Pylkkänen, 2024; Krogh & Pylkkänen, 2024).

### Procedure

Prior to MEG recording, each participant’s head shape was digitally recorded using a Polhemus FastSCAN device. These three-dimensional models were used to constrain the source localization models computed in later analysis stages. Before the MEG recording, each participant performed 10 practice trials. Afterwards, they were escorted into a dimly lit magnetically shielded room where they performed the entire experiment while in the supine position with their MEG recorded. The experiment was presented using the PsychoPy package (Peirce et al., 2010) in Python roughly 50 cm away from the participant’s face. Stimuli were presented using a white Courier New font with a visual angle of 0° 34’ against a grey background. Trials were presented to each participant in a random order, and the participants had the option of taking a break every time they completed a tenth of the trials. Once participants began the experiment within the magnetically shielded room, the sessions lasted roughly one hour. In addition to this session, participants were recruited to participate in a separate experiment using different stimuli that addressed a question other than the one addressed in this study.

### MEG data acquisition and preprocessing

MEG data were collected using a 157-channel axial gradiometer system (Kanazawa Institute of Technology, Tokyo, Japan) with a sampling rate of 1000 Hz. During acquisition, a high-pass filter at 1 Hz was applied to reduce environmental noise, along with a low-pass filter at 200 Hz. Following data collection, noise reduction was performed using the continuously adjusted least-squares method (Adachi et al., 2001) within the MEG 160 software (Meg Laboratory 2.004A, Yokogawa Electric Corporation, Kanazawa Institute of Technology). Additional low-pass filtering at 40 Hz was subsequently implemented using MNE-Python (Gramfort et al., 2013; 2014). Channels exhibiting irregularities upon visual inspection were excluded, and independent component analysis (ICA) was utilized to identify and remove artifacts related to eye blinks and cardiac activity. The preprocessed data were then segmented into epochs spanning from 100 ms before the onset of Sentence One to 800 ms after, resulting in 900 ms epochs. The first 100 ms of each epoch served as a baseline for computing the noise-covariance matrix. Epochs with sensor values exceeding 3000 fT at any point were rejected.

Source-level activity was estimated from the evoked responses using dynamical statistical parameter mapping (dSPM) (Dale et al., 2000). Each participant’s head shape and fiducial landmarks, recorded prior to the experiment, were used for alignment and morphing onto the “fsaverage” brain model via FreeSurfer software (http://surfer.nmr.mgh.harvard.edu/). MEG activity across conditions was averaged, and the forward model was computed using the Boundary Element Model (Bonnet, 1996; Mosher et al., 1992). Covariance matrices were derived from the 100 ms preceding stimulus onset. The source-level inverse solution and neural activity estimates were obtained using minimum norm estimation (Hämäläinen and Ilmoniemi, 1994).

### Behavioral data analyses

Behavioral data were collected to examine the presence of a behavioral sentence superiority effect (SSE). This effect consists of reduced reaction times and/or higher accuracies for fully grammatical sentences relative to non-compositional stimuli and has been in observed in many studies employing parallel presentation (Snell and Grainger, 2017; Massol et al., 2021; Vandendaele and Grainger, 2025). Data cleaning involved excluding trials with reaction times (RTs) shorter than 200 ms and longer than 4000 ms. Next, RTs were standardized within each participant by computing z-scores, and trials falling more than three standard deviations from a participant’s mean RT were removed. Separate models were constructed for RT and accuracy, analyzing match and mismatch trials independently, as these conditions likely impose distinct cognitive demands. Reaction times were analyzed using linear mixed-effects models, with composability and stimulus size as fixed effects and subject as a random effect. Attempts to include items as a random effect resulted in convergence issues. Accuracy data were analyzed using a generalized linear mixed-effects logistic regression model with the same predictors and structure as the RT models. Pairwise comparisons were conducted via likelihood ratio tests and adjusted using Tukey’s correction. All behavioral analyses were performed using the lme4 (Bates et al., 2015) and afex (Singmann et al., 2023) packages in R (v4.3.1).

### MEG data analyses

#### ROI analysis

We performed cluster-based permutation tests (Maris and Oostenveld, 2007) across the left hemisphere Brodmann Areas (BAs) 11, 20, 21, 22, and 38, given the literature implicating these regions in lexical and compositional processing (Pylkkänen, 2019; Matchin and Hickok, 2020; Flick and Pylkkänen, 2021). Within each region of interest (ROI), the sources were mean flipped at each time point for each condition. The resulting time courses were then fed into the temporal cluster-based permutation test, with an analysis window of 0 – 800 ms after stimulus onset, cluster-forming threshold of p < .05, 10,000 permutations to determine corrected cluster p-values (alpha < .05), and finally, FDR correction across ROIs. The point-by-point statistic to derive temporal clusters was an F-test following a 3 x 4 repeated measures ANOVA design, with the factors stimulus size (small, medium, large) and determiner *(all*, *some*, *no*, *the).* The three levels of the size factor consisted of bare determiners, determiner phrases, and sentences. The determiner factor was included to investigate topics separate from the current study.

The goal of our ROI analysis was to identify effects of composition, which were defined in the following way. We first performed the repeated measures ANOVA using the size factor over the averaged source estimates with the time window being 0 – 800 ms. Then, for each significant cluster revealed by the analysis, we performed pairwise comparisons between the non-compositional list conditions using neural activation averaged within the significant time window. We interpret clusters with *no* significant comparisons among the list conditions as reflecting *combinatory* processing. In contrast, clusters with *any* observed significant comparisons among the list conditions was considered to reflect aspects of processing shared by both types of stimuli, such as the detection of form properties, namely the size of the stimulus.

Our hypotheses differ with respect to the expected number of observed combinatory processing stages. GLOSA and GLOPA predict size effects followed by combinatorial effects. Size effects correspond to the “snapshot” processing posited in both hypotheses. On the other hand, GLU predicts to see overlapping stages of combinatory processing with the encoding of the stimulus.

#### Generalized Linear Model (GLM) analysis

For each significant effect of size, whether specific to the combinatory stimuli or not, a two-stage generalized linear model (GLM) analysis within the effect’s time window (plus and minus 25 ms) was conducted to probe the effect’s functional profile. Specifically, we assessed whether the significant cluster showed hallmarks of lexical or compositional processing, operationalized by lexical frequency and bigram frequency respectively. We fitted two classes of models with the single-trial source estimates as dependent variables. The first class modeled lexical frequencies and the second modeled bigram frequencies. Lexical frequency and bigram frequency were computed using the Corpus of Contemporary American English or *Coca* (Davies, 2008). For the compositional stimuli, we ran both models on the determiner phrases and the sentences. We did not run lexical frequency models on the bare determiners, given the relative lack of variance in lexical frequency within these conditions relative to others. For the determiner phrases, we computed the lexical frequency model *dspm ∼ log_frequency(1) + log_frequency(2)* and the bigram frequency model *dspm ∼ bigram(1)* and for the sentences, the lexical frequency model *dspm ∼ log_frequency(1) + log_frequency(2) + log_frequency(4)* and the bigram model *dspm ∼ log_bigram(1) + log_bigram(2) + log_bigram(3).* The lexical frequency model for the sentences did not include the logged frequency for the third word, because this word did not vary at all within our stimuli. For the non-compositional stimuli, we computed only lexical frequency models, given that the bigram frequencies of adjacent plural nouns were uniformly zero or close to zero in COCA. Thus, we only computed models for lexical frequency following the same pattern as above for the phrases and sentences.

At the first stage, for each subject, we fitted a linear regression model to the appropriate subset of the subject’s data across every source point. For each predictor in the linear model, this yielded a set of 5124 time-courses corresponding to the beta coefficient of that predictor at every time point in the epoch. At the second stage, we performed one-sample spatiotemporal clustering tests to examine the functional properties of the Brodmann Areas showing significant effects for size in the repeated measures ANOVA tests conducted above. For each BA showing a significant size effect, we ran a two-tailed spatiotemporal clustering test using the BA as a search window and the time window of the significant effect buffered with 25 ms before and after. We used a p-value threshold of less than 0.05, and corrected cluster p-values were estimated with 10,000 permutations. These analyses were carried out using the eelbrain package in Python (Brodbeck et al., 2023).

## RESULTS

### Behavioral results

Our test stimuli were followed by a second word string that was either identical to or differed by one word from the critical stimulus. Subjects (N = 24) judged whether the strings matched, as in prior rapid parallel visual presentation (RPVP) studies (Flower and Pylkkänen, 2024, Fallon and Pylkkänen, 2024). Overall, match trials (795.1 ms ± 396.1 ms) were faster than mismatch trials (844.0 ms ± 401.3 ms), and match accuracy (96.12% ± 19.31%) slightly exceeded mismatch accuracy (93.99% ± 23.78%) (Fig. 3).

**Figure 2:**
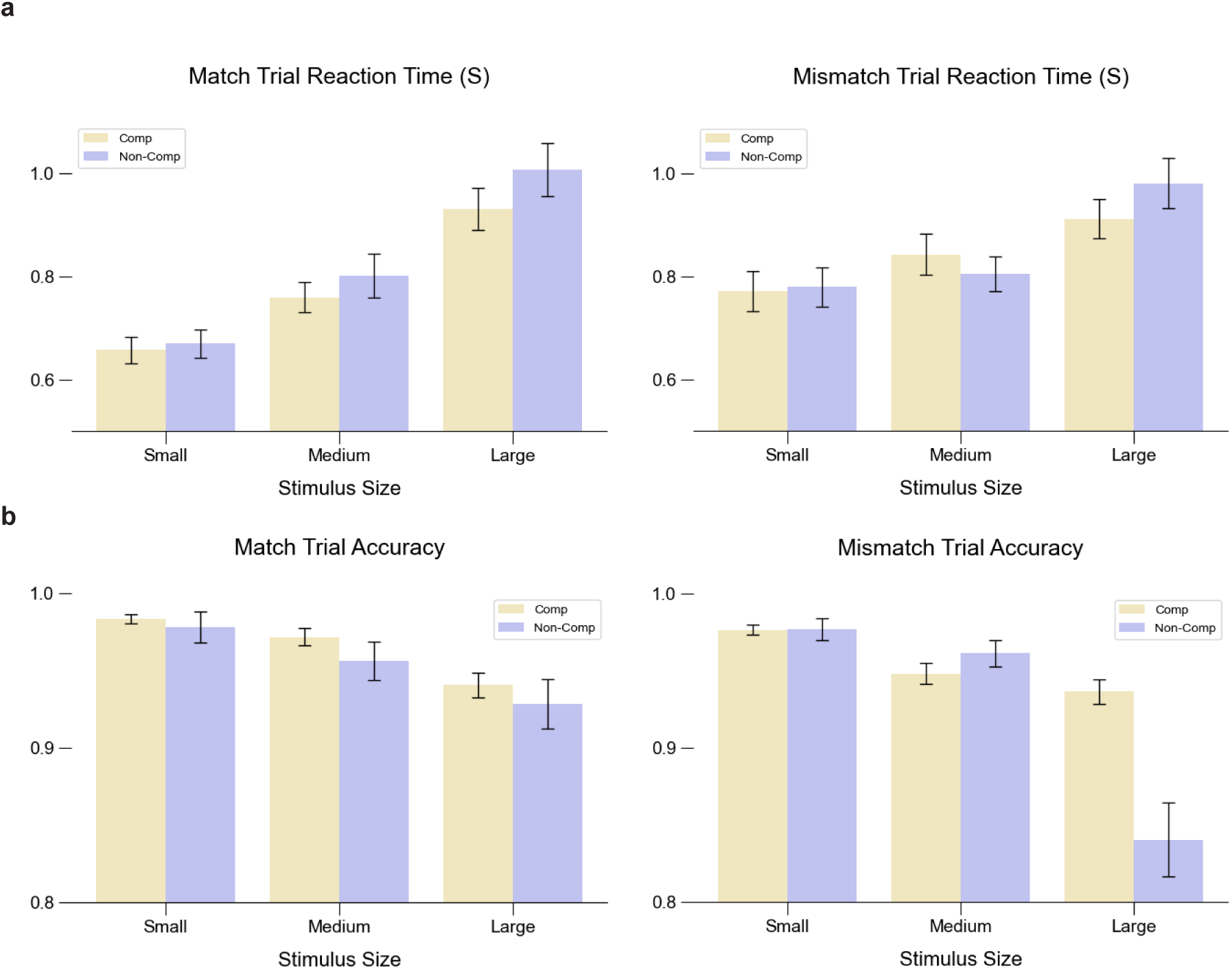
RT and accuracies show a clear behavioral cost for increasing stimulus sizes. (A) Within both match and mismatch trials sentences (comp, large) elicited significantly longer RTs than noun-lists (large, non-comp). (B) Similarly, both match and mismatch trials show a behavioral cost for noun-list stimuli relative to full sentences.

#### Behavioral Instances of SSE and PSE

Pairwise comparisons between the sentences and the three-noun lists showed greater RTs for the lists relative to the sentences in both Match (*p* = 0.0088) and Mismatch trials (*p* = 0.0021), providing evidence of an SSE. We further find evidence of an SSE in higher accuracies for sentences relative to lists in the Match (*p* = 0.0453) and Mismatch trials (*p* = 0.0001). Although these conditions differ in total number of words, this effect cannot be reduced to the sentences having an extra word, because we would expect the three-noun lists to elicit quicker RTs. Thus, it is safe to conclude that the increased RTs for both match and mismatch trials in this comparison is due to the sentences consisting of grammatical strings of English, while the noun lists are not composable, consistent with much previous work on the SSE (Snell and Grainger, 2017; Massol et al., 2021; Flower and Pylkkänen, 2024; Fallon and Pylkkänen, 2024; Vandendaele et al., 2025). We further find evidence of a behavioral *phrasal superiority effect* (PSE) in match trial RT (*p* = 0.0142) and a trending effect in mismatch RT (*p* = 0.054). However, there was no evidence for a PSE in the accuracies (Match: *p* = 0.47; Mismatch: *p* = 0.6). This is a novel finding, given the absence of PSEs reported in the parallel presentation literature.

#### Main Effects of Stimulus Size

As expected, we observe a consistent effect of stimulus size for RTs across all comparisons for both match and mismatch trials. Within the match trials, we find that RTs for large stimuli (949.8 ms ± 441.2 ms) were greater than those of medium stimuli (774.4 ms ± 371.0; *p* < 0.0001), which were in turn greater than those of small stimuli (668.4 ms ± 318.6 ms; *p* < 0.0001). The mismatch trials exhibit the same pattern of large stimuli (932.7 ms ± 407.5 ms; *p* < 0.0001) eliciting higher RTs than the medium-sized stimuli (834.6 ms ± 398.9 ms) followed by the small stimuli (766.9 ms ± 380.3 ms; *p* < 0.0001). Match trial accuracy for large stimuli (94.5% ± 22.8%) was not significantly different from medium-sized stimuli (95.9% ± 19.8%; *p* = 0.2729), but accuracy for small stimuli (97.9% ± 14.5%) was significantly higher than both the large (*p* = 0.0002) and medium-sized conditions (*p* = 0.022). For the mismatch trials, large stimuli had significantly worse accuracies (88.6% ± 31.7%) than both small (97.5% ± 15.7%; *p* < 0.0001) and medium-sized stimuli (95.8% ± 20.2%; *p* < 0.0001), but there was no difference between the small and medium-sized stimuli (*p* = 0.94).

### MEG results

#### ROI Results

For each region, we observed two distinct stages of processing, not a set of overlapping stages. Each region showed either two stages of size processing between combinatory stimuli and lists, or one stage of size processing followed by a stage of combinatory processing. The observed sequence of size processing followed by combinatory processing is consistent with the GLOSA and GLOPA hypotheses. The regions exhibiting this behavior were BA 20, BA 21, and BA 38, whereas BA 11 and BA 22 showed evidence for two stages of size processing. We first present the results from regions exhibiting size effects followed combinatory effects (ordered by latency of the combinatory processing stage) followed by the regions that showed two stages of size processing.

#### BA 21

In BA 21, we observed two significant clusters, with the first (119 – 197 ms, *p* = 0.008) showing a profile of size processing and the second (291 – 403 ms, p = 0.0001) a profile of combinatory processing (Fig. 4). The first cluster exhibited a three-way contrast among the compositional conditions with sentences eliciting increased activation relative to bare determiners (*p* = 0.005) and phrases (*p* = 0.003), and phrases eliciting increased signals relative to bare determiners (*p* = 0.5). Further, the comparisons in the non-compositional stimuli showed a significant increase of activity for the three-noun lists relative to the single nouns (*p* = 0.03), but not for other comparisons. The sensitivity to the non-combinatory stimuli thus qualifies this cluster as reflecting size processing.

Within the latter cluster, sentences elicited significantly greater activity compared to bare determiners (*p* < 0.0001) and phrases (*p* < 0.0001), but phrases and determiners did not yield any significant differences. Comparisons between the non-compositional stimuli did not yield any significant differences between conditions. Thus, the neural activity within this cluster shows sensitivity to combinatory processing that cannot be simply explained by differences in the size of the stimuli. Similarly with the following two ROIs, the lack of multiple stages of combinatory processing are consistent with GLOSA.

**Figure 3:**
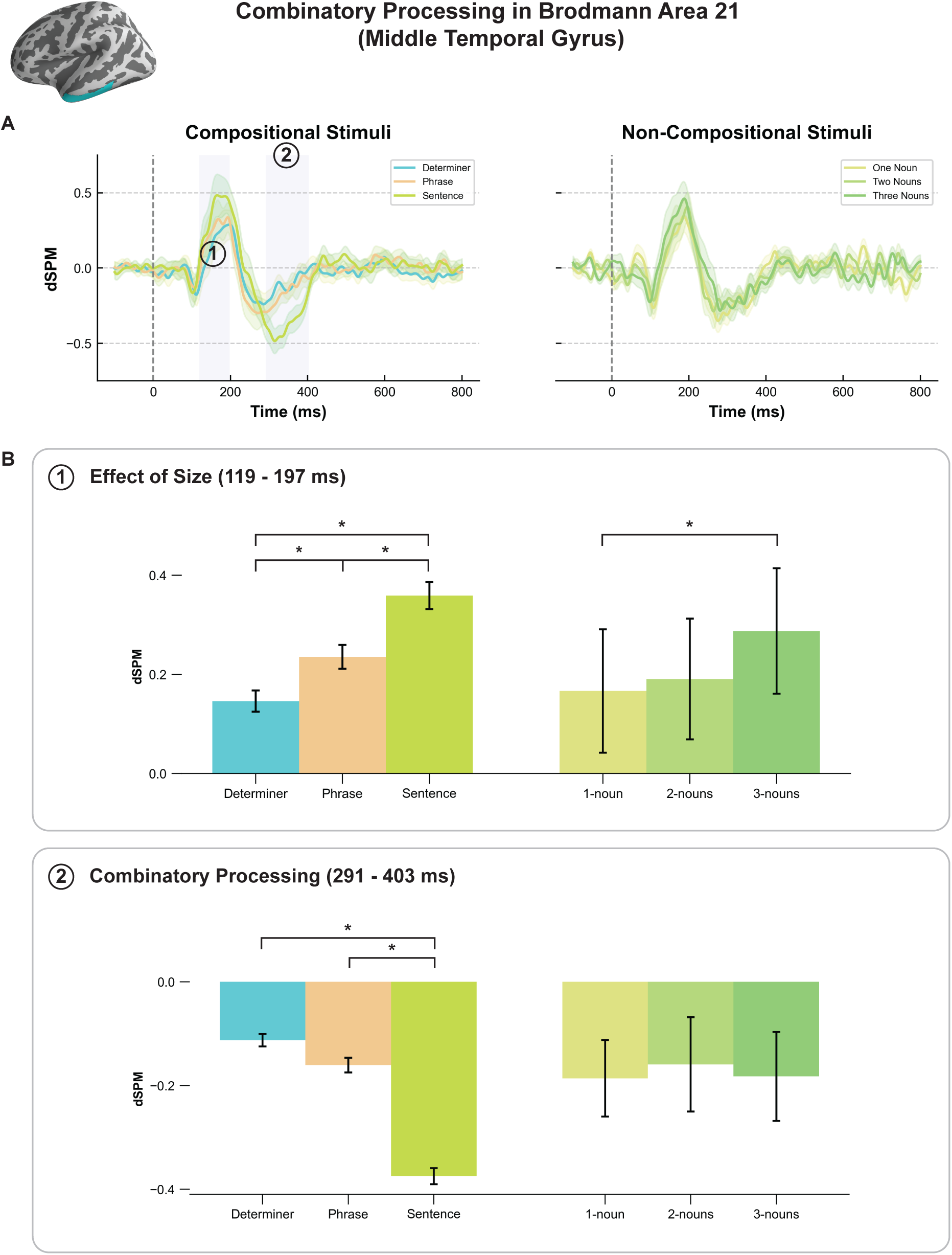
Combinatory processing in MTG. (A) Temporal clustering test using factor of size reveals two clusters of activity (left). The first cluster takes place 119 – 197 ms after stimulus onset. The second cluster takes place 291 – 403 ms after stimulus onset. (B) Pairwise comparisons showed evidence for size processing in the early cluster and combinatory processing in the late cluster. This pattern of effects is consistent with single stage processing.

#### BA 38

BA 38 also showed an early cluster (125 – 255 ms, *p* = 0.0002) with a size processing profile and a later one (298 – 440 ms, *p* = 0.0001) with a combinatory one (Fig. 5). In the first cluster, pairwise comparisons found that the larger stimuli (sentences and three-noun lists) elicited significantly greater negative amplitudes than smaller sized stimuli (sentence vs. determiner: *p* = 0.0002; sentence vs. phrase: *p* < 0.0001; three nouns vs. one noun: *p* = 0.0001; three nouns vs. two nouns: *p* = 0.003). There were no significant differences between the bare determiners and the determiner phrases; however, there was a trending effect for the comparison between single nouns and the two-noun conditions (p = 0.055). Thus, this effect showed increased negative amplitudes for large-sized stimuli relative to smaller stimuli, consistent with size processing. Within the second cluster, pairwise comparisons showed significantly greater activity for determiner phrases (*p* = 0.04) and sentences (*p* < 0.0001) relative to bare determiners as well as between phrases and sentences (*p* < 0.0001). No comparisons were found among the non-combinatory stimuli, thus demonstrating combinatory processing behavior.

**Figure 4:**
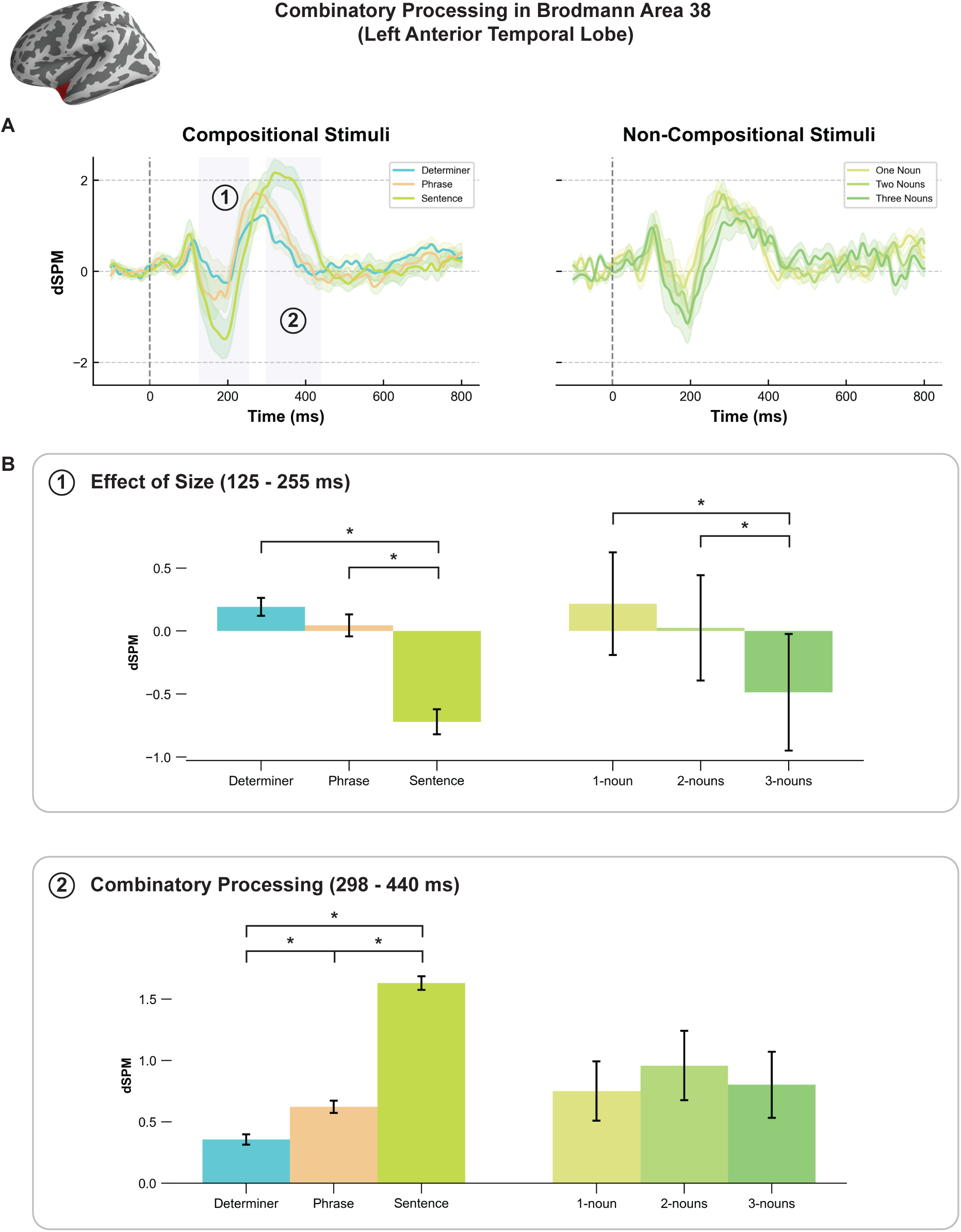
Evidence for GLOSA in LATL. (A) Temporal clustering test using factor of size found early cluster between 125 – 255 ms, and a later cluster 298 – 440 ms after stimulus onset. (B) Pairwise comparisons showed evidence for size processing in first cluster and combinatory processing in the second cluster, thus evidence for the LATL performing single stage processing.

#### BA 20

Finally, the same pattern was seen in BA 20, with an early cluster (125 – 253 ms, *p* = 0.001) showing evidence of size processing followed by a later one (510 – 576 ms, *p* = 0.04) that patterned according to combinatory processing (Fig. 6). The early effect could be described as primarily reflecting a main effect of size despite the lack of significant difference in activation between the determiners and the determiner phrases. Within the compositional conditions, the sentences elicited increased activity relative to bare determiners (*p* = 0.0002) and determiner phrases (*p* = 0.001). However, there was a three-way distinction between the one, two, and three-noun stimuli, where three-noun lists elicited increased activity compared to two-noun lists (*p* = 0.002) and single nouns (*p* < 0.001), and two-noun lists elicited increased activity compared to single nouns (*p* = 0.03). The activity in this cluster is thus reflective of size processing. The second cluster exhibited compositional behavior; both sentences (*p* = 0.01) and phrases (*p* = 0.0009) yielded greater activation relative to bare determiners, but there was no significant difference between the sentence and phrase conditions. On the other hand, there were no significant differences between the non-compositional stimuli, which is consistent with combinatory processing.

**Figure 5:**
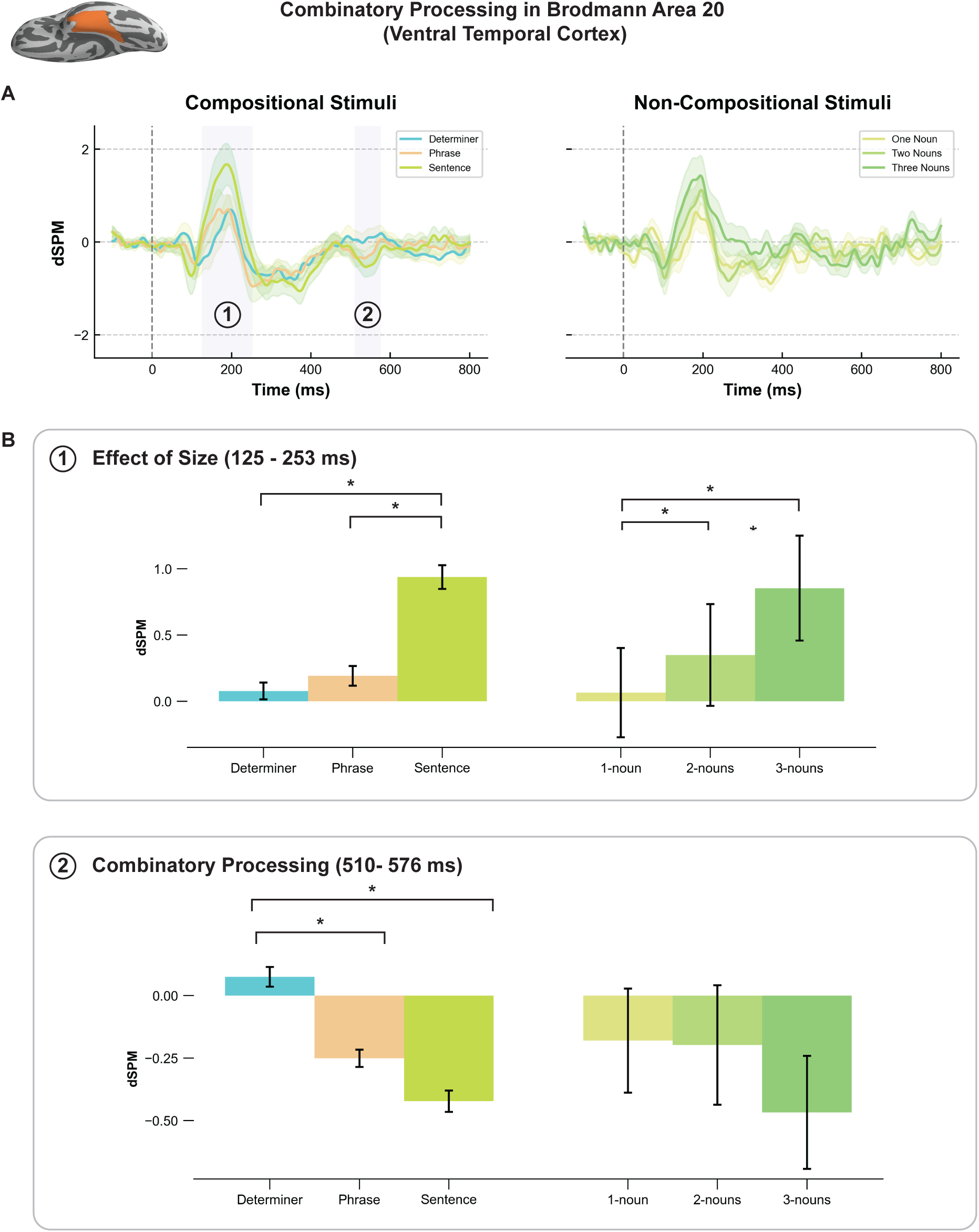
Combinatory processing in ventrotemporal cortex. (A) Early cluster was found 125-253 ms after stimulus onset, and a later cluster was found 510 – 576 ms after stimulus onset. (B) Pairwise comparisons showed evidence for size processing in first cluster and combinatory processing in the second cluster. This is consistent with single stage processing.

#### BA 11

Finally, our ROI analysis found two clusters in BA 11 showing size processing behavior (Fig. 7a). In the first cluster (116 – 250 ms, *p* = 0.0003), Pairwise comparisons in this cluster showed a three-way difference in neural activation across the different structures. Sentences elicited increased activity compared to phrases (*p* < 0.0001) and bare determiners (*p* < 0.0001), and phrases elicited increased activity relative to bare determiners as well (*p* = 0.03). Comparisons in the non-compositional conditions showed a similar, but not identical pattern. Three-noun lists elicited more activity than two-noun lists (*p* = 0.02) and single nouns (*p* = 0.0009). In the later cluster (300 – 446 ms, *p* = 0.0001), activity among compositional stimuli showed a three-way contrast across structures. Sentences elicited increased negative activity relative to phrases (*p* < 0.0001) and bare determiners (*p* < 0.0001), and phrases elicited increased negative activity relative to bare determiners (*p* = 0.02). Comparisons showed a difference between only the two-noun lists and the single noun conditions (*p* = 0.05) and no other significant comparisons.

**Figure 6:**
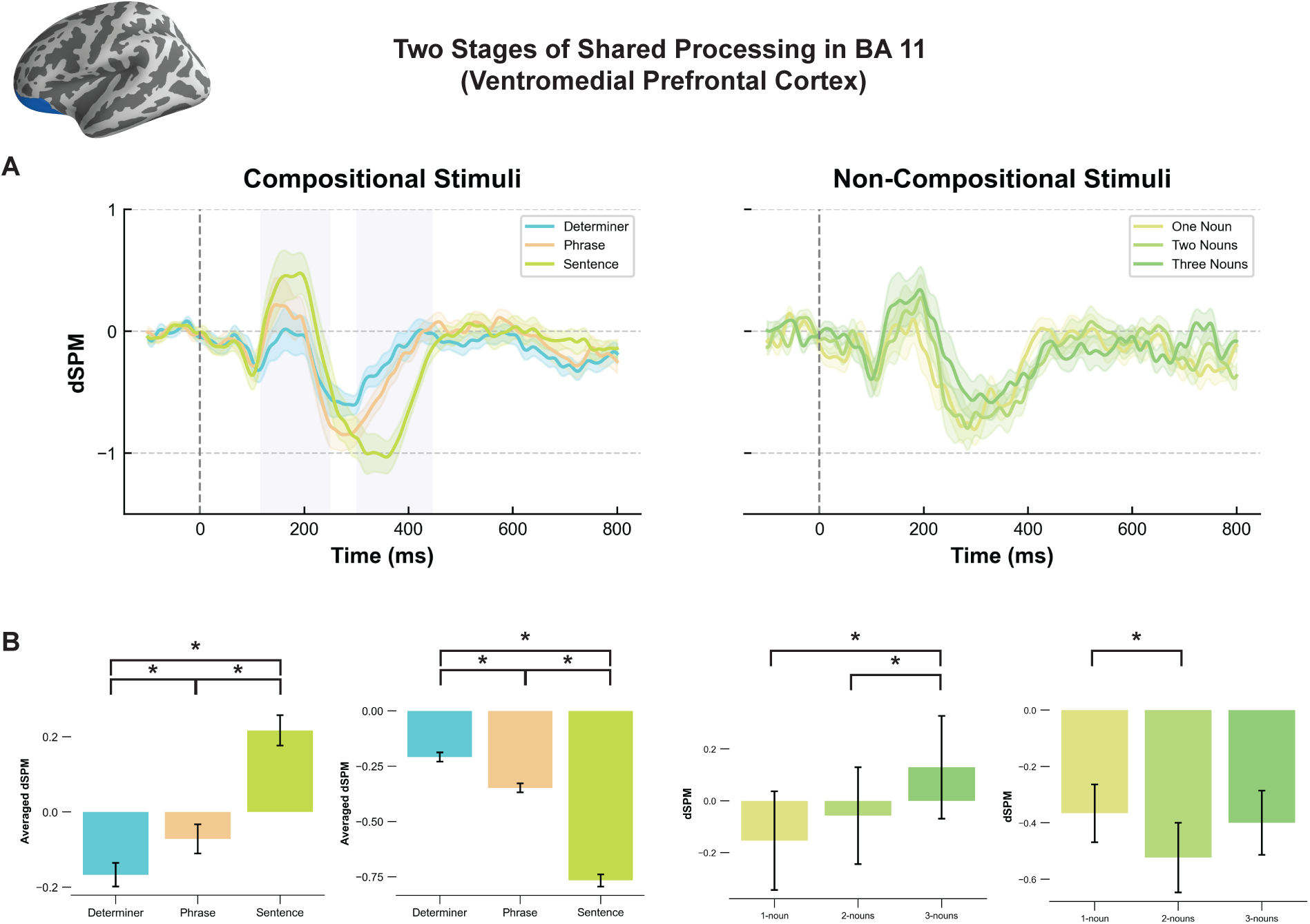
Two stages of size processing in vmPFC. (A) Within BA 11, an early cluster was found 116 – 250 ms followed by a later cluster 300 – 446 ms (B) Both clusters show sensitivity to the size of the stimulus regardless of composability.

#### BA 22

Unlike the previous three ROIs, our ROI analysis revealed two clusters in BA 22 that only reflected size processing (Fig. 7b). Pairwise comparisons in the first cluster (107 – 160 ms, *p* = 0.0457) revealed significantly higher activity for sentences relative to bare determiners (*p* = 0.0027) and phrases (*p* = 0.01) but no significant differences between phrases and sentences. Comparisons within the non-compositional conditions showed significantly more activity for three-noun lists and single nouns only (p = 0.0006), thus displaying size processing. The second cluster (332 – 391 ms, *p* = 0.002) exhibited a pattern of activity similar to the first cluster, where sentences elicited more activity than bare determiners (*p* = 0.002) and determiner phrases (*p* < 0.0001) but no significant differences between phrases and sentences. Again, the non-compositional stimuli also showed significantly more activity for three-noun lists compared to single nouns only (*p* = 0.02), consistent with size processing.

**Figure 7:**
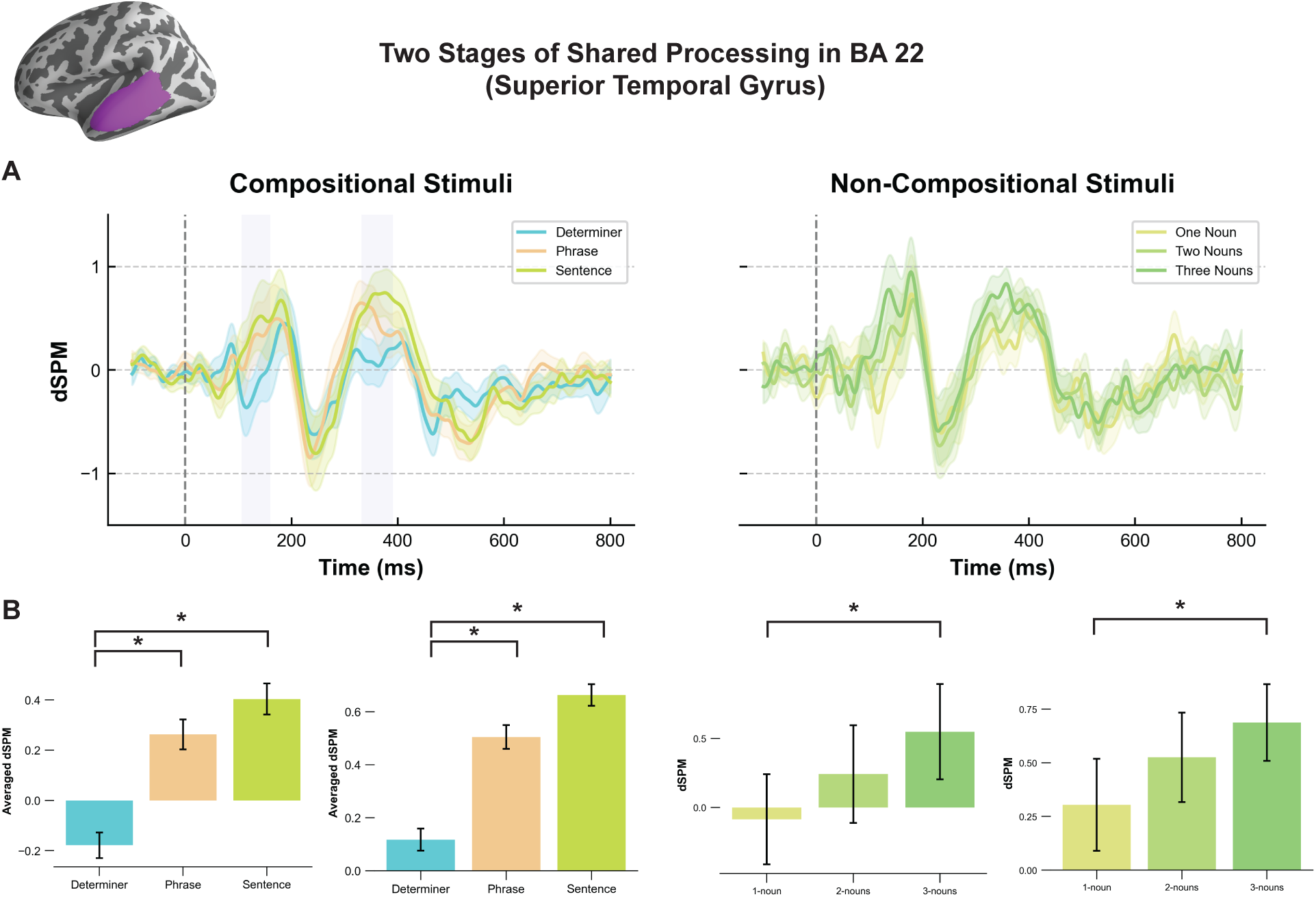
Two stages of size processing in STG. (A) Within BA 22, an early cluster was found 107 – 160 ms followed by a later cluster 332 – 391 ms. (B) Both clusters show sensitivity to the size of the stimulus regardless of composability.

#### GLM results

The ROI analysis showed that all five ROIs showed effects of size. Thus, to test for evidence of serial or parallel processing in each of these regions, we ran GLM models of lexical and bigram frequency within each of the ROIs using the time windows of the significant clusters padded by 25 ms of activity before and after. The GLM analysis revealed *serial* effects of bigram frequency in BA 38, which is consistent with GLOSA. Within the sentences, BA 38 showed an effect for the first bigram, “all cats,” at 231-332 ms after stimulus onset (*p* = 0.02) and to the second bigram, “cats are,” at 372-444 ms after stimulus onset (*p* = 0.01). Interestingly, for the phrases, BA 38 showed sensitivity to the first (and only) bigram from 212 to 333 ms after stimulus onset (search window: 125 – 440 ms; *p* = 0.01). That is to say, sensitivity to the bigram “the cats” when presented alone occurred roughly 20 ms before the bigram “the cats” when presented in a full sentence such as “the cats are nice.” No other effects were found in other ROIs that served to distinguish between serial and parallel processing hypotheses. We present the other results from the GLM analysis starting with the lexical frequency models.

The linear models using lexical frequency as predictors showed significant effects in only BA 22. Within BA 22 (search window: 307 – 416 ms), the analysis revealed a significant effect for only the second word at 328-375 ms (p = 0.02), and this effect surfaced when looking at the sentence conditions. No significant effects were found for lexical frequency in any other region or other conditions. The timing of this effect is consistent with studies finding effects of lexical access for single words taking place within the superior temporal gyrus (STG) during this time window (Pylkkänen et al., 2002; Pylkkänen and Marantz, 2003, Lau et al., 2008).

The bigram frequency models further showed effects within BA 11 and BA 21. These effects are expected given that these regions are broadly implicated in a network of cortical regions engaging in linguistic combinatorics (Pylkkänen, 2019). There was also a significant effect for the first bigram in BA 11 from 233 to 358 ms after stimulus onset (search window: 116 – 446 ms; *p* = 0.01). Finally, BA 21 also exhibited a significant effect of bigram frequency for the second bigram, “cats are” between 373 and 428 ms (search window: 266 – 428; *p* = 0.01). These bigram frequency effects are consistent with an extensive literature implicating these regions in compositional linguistic processing.

**Figure 8:**
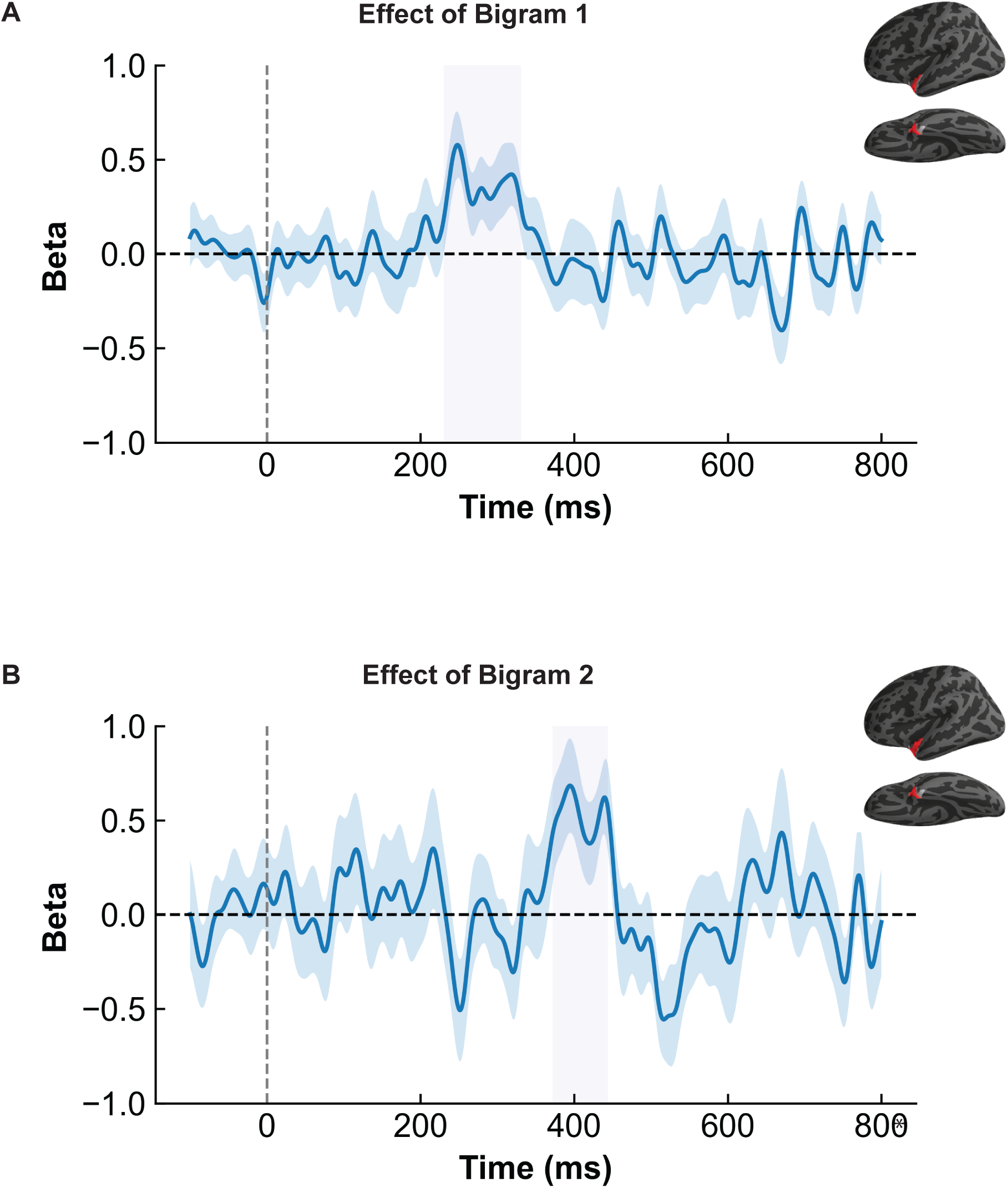
Serial processing in the combinatory stage of BA 38. For sentences, GLM analyses revealed sequential sensitivity to the first two bigrams (*the cats* and *cats are*). The first bigram showed sensitivity 212 – 333 ms after stimulus onset, and the second bigram showed sensitivity 372 – 444 ms after stimulus onset.

## DISCUSSION

In this work, we sought to understand the relationship between the brain’s evoked response to a full sentence as compared to a single word, a question tightly linked with the degree of serial versus parallel processing when comprehending complex linguistic stimulus. Our results compellingly show that evoked responses for sentences are indeed highly similar to those of single words, with each effect manifesting as a modulation of a shared waveform, as opposed to sentences and single words eliciting qualitatively different responses. Within this shared response pattern, our analyses directly examined when, after stimulus onset, the combinatory properties of a stimulus begin to influence neural activity. A set of ROIs considered combinatory in a sizeable prior literature—i.e., the LATL (BA 38), MTG (BA 21), ventral temporal cortex (BA 20), and possibly the ventromedial prefrontal (BA 11) cortex—all showed a two-stage behavior, with early bottom-up encoding of stimulus size followed combinatory processing in a single stage, consistent with GLOSA. Finally, within the LATL, we also observed evidence of serial left-to-right composition, as predicted by GLOSA.

### Similarity of Evoked Responses to Words and Sentences

We find that the evoked responses to sentences and words are not fundamentally different, as already suggested by the patterns of evoked responses in prior studies, although they did not yet include direct comparisons between single words and multi-word stimuli (Wen et al., 2019; 2021). In each of our ROIs, all our size and composition effect were modulations of a waveform that had the same general shape for single words and larger expressions. This outcome is inconsistent with the fully serial and GLU hypotheses. Prior work has already shown that the N400, classically associated with lexical expectancy, is likewise sensitive to the well-formedness of full sentences Wen et al. (2019; 2021), suggesting a shared processing profile for words and sentences. This hypothesis also has a strong predecessor in hemodynamic literature, motivated by the shared sensitivity of language regions to both lexico-semantic and syntactic processing (Fedorenko et al. 2020; Shain et al., 2024). In theoretical linguistics, the view that word and sentence level structure are governed by the same syntactic principles has been influential for decades (Halle and Marantz, 1993; Starke, 2009). Experimental work on language has, however, been dominated by the so-called “lexicalist” notion that words and sentences are representationally distinct, as recently discussed and critiqued by Krauska and Lau (2023), advocating for a move away from lexicalist assumptions. Thus, a convergence may be arising that the that the conventional theory positing a fundamental difference between words and sentences has run its course both as a model of linguistic representation as well as of processing.

### GLOSA Processing in the LATL

The profile of the LATL followed the GLOSA hypothesis: an initial “snapshot” stage of size effects, shared between combinatory and noncombinatorial stimuli, was followed by a stage of composition effects showing evidence of serial processing. Specifically, in the composition stage, the frequency of the first bigram (*all cats*) impacted LATL signals at 212 – 333 ms and that of the second bigram (*cats are*) at 372 – 444 ms after stimulus onset. We did not observe effects of third bigram frequency, possibly because of English readers’ leftward bias (Reichle et al., 2003). Since readers of Roman scripts preferentially deploy attention to the left-hand side of stimuli, one might reasonably expect a higher amount of variance in neural activity devoted to encoding properties of the right side of a large stimulus such as a four-word sentence. This can be seen in other studies using parallel presentation that show slightly reduced accuracies for words on the right edge of a sentence compared to the left edge (Snell and Grainger, 2017).

These results speak to a longstanding debate within the psychology of reading concerning whether readers are capable of semantically processing multiple words simultaneously. On the one hand, serial models of reading propose that the mind is only capable of semantically processing a single word at a time (Reichle et al., 1998; 2003; 2009). According to these accounts, lexical representations are activated one at a time, and this is due to a processing bottleneck that prevents extraction of meaning from multiple words at once. The opposing view is that lexical representations can be accessed in parallel, although details concerning combinatory processing are usually not discussed (Snell and Grainger, 2019). In this kind of model, the processing principles that underlie visual word recognition are simply scaled up to the recognition of full sentences. For words, sets of letter detector nodes activate candidate word nodes, which then feedback activation to the letter detector nodes, eventually resulting in a single candidate word node reaching an activation threshold. When applied to full sentences, it is word nodes that are initially activated, feeding into nodes for full sentences, etc.

The LATL’s early sensitivity to the first bigram shows that as early as 230 ms after a full sentence is presented, representations that encode word-level features of the sentence’s first two words have already been activated. However, the results do not suggest that the brain’s capacity for activating higher-level combinatory information of bigrams is unlimited. The serial combinatory effects that we see in the LATL instead suggest that processing may be limited to a single bigram at a time. Thus, the processing bottleneck might not be over words themselves but over higher-order combinable representations. This is consistent with Anupindi et al. (2025) who find that under highly constrained presentation times, readers are only capable of recognizing two words at once if they form a compound. In other words, meaningful bigrams may be recognized in parallel, but not two nouns that do not form a single combinable unit.

### Combinatory Profiles of the MTG and vmPFC

In the MTG, we observed an early stage of size-related activity followed by a later stage combinatory processing, aligning with the snapshot-then-composition sequence predicted by the GLOSA and GLOPA hypotheses. However, neither lexical nor bigram frequency analyses yielded effects that determined whether the observed combinatory activity reflects serial or parallel integration. Accordingly, these results are inconsistent with GLU but cannot distinguish between GLOSA or GLOPA.

Our MTG findings align with a growing body of MEG evidence implicating this region in combinatory, and specifically syntactic, processing (Matchin et al., 2019; Flick and Pylkkänen, 2020; Matar et al., 2021; Law and Pylkkänen, 2021; Fallon and Pylkkänen, 2024). Computational modeling studies that link neural responses with syntactic parsing operations have likewise identified the MTG as a locus of syntactic composition (Brennan et al., 2016; Brennen et al., 2020; Stanojević et al., 2023; Sugimoto et al., 2024; Coopmans et al., 2025). The temporal profile of our combinatory response overlaps with the N400 component (Halgren et al., 2002), traditionally associated with lexical-semantic access (Lau et al., 2008). However, this interpretation is unlikely here, as we found no differences among non-combinatory stimuli of varying sizes. Instead, our results are consistent with proposals that the MTG supports lexicalized combinatory processing—a mechanism sensitive to both lexical and syntactic properties during composition (Matchin et al., 2019; Matchin and Hickok, 2020; Shain et al., 2024).

It is noteworthy that our GLM analysis did not reveal any significant effects of lexical frequency within the MTG, despite numerous reports of such effects in single-word and serial presentation paradigms (Lau et al., 2008; Fruchter et al., 2015; Huizeling et al., 2022). In contrast, no prior parallel presentation studies have identified lexical frequency effects in this region, and only one EEG study reported frequency sensitivity under parallel presentation (Dunagan et al., 2025), though the neural sources of that effect remain uncertain given EEG’s limited spatial resolution. The absence of lexical frequency effects in the present study may therefore reflect factors specific to our stimulus set and modeling approach. First, the range of lexical and bigram frequencies in our materials was likely too restricted to elicit measurable frequency-dependent responses, as the stimuli were designed for other experimental goals. In particular, the repeated use of a small determiner set (“all,” “some,” “no,” “the”) constrained the variability of early lexical and combinatory contexts. Second, our frequency estimates were derived from surface forms rather than lemmas. Because our noun stimuli were pluralized (e.g., cats), lemma-based frequencies may have provided a more accurate reflection of lexical access processes in the brain. Finally, a two-stage snapshot-then-composition profile was also seen in BA 11, the ventromedial prefrontal cortex (vmPFC). Here the evidence for the composition specificity of the second stage was slightly less crisp than for BAs 38 or 21, given that two-word lists did trend (*p* = .05) towards larger amplitudes than single words. However, this trend did not continue on to three-word lists, which diverged neither from one nor two-word lists. Thus overall, the pattern was still clearly different from the combinatory stimuli, for which the vmPFC signals robustly tracked the number of words and thus the amount of composition. Like BA21, this activity was insensitive to frequency. Along with the LATL, the vmPFC has shown effects of composition in a large number of MEG studies (Pylkkänen, 2019), including both comprehension and production (Bemis & Pylkkänen, 2011; Pylkkänen et al., 2014), simple and complex composition (Pylkkänen & McElree, 2007) as well as controlled and naturalistic studies (Brennan & Pylkkänen, 2012). In prior comprehension studies using serial presentation, the timing of vmPFC activation has been highly consistent with what is observed here for parallel presentation, around 400 ms. Thus, even though in this study the brain encountered all combining words simultaneously, this did not delay the vmPFC combinatory signals, an outcome that would be unexpected under a theory requiring words to be retrieved sequentially before combination.

## CONCLUSION

This study was inspired by the observation that full sentences presented in parallel elicit electrophysiological responses that look remarkably similar to those well established for single words (Wen et al., 2019, 2021). This raises the question of whether words, phrases, and sentences all elicit waveforms with similar morphology, passing through a pathway in which the same form, structure, and meaning detection mechanisms operate for any linguistic input. Under theories in which all words are structured entities like phrases and sentences, such a processing account is natural (Halle & Marantz, 1993; Marantz, 1997). To assess this within subjects, we presented single words, phrases, and sentences in parallel to observe whether the waveform morphologies for these three stimulus types are shaped similarly. The answer was a resounding yes: across all brain regions examined, each area elicited the same sequence of peaks and valleys for single words and syntactically complex stimuli. Within these responses, we assessed which components showed sensitivity to composition that could not be explained by the sheer size contrast between single words, phrases, and sentences. The left anterior and posterior temporal cortices, as well as the vmPFC, all showed such responses. Of these, the signals in the left anterior temporal lobe were frequency-modulated in a way that suggests a serial, left-to-right processing mechanism. Thus, overall, our findings support a Global-to-Serial Assembly (GLOSA) model in which the brain first detects the global form of the stimulus in a snapshot-like manner and then probes its combinatory properties, with at least some serial left-to-right dynamics.

## ACKNOWLEDGEMENTS

This work was supported by the National Science Foundation (BCS-2335767) and by award G1001 from NYUAD Institute, New York University Abu Dhabi (LP).

